## Supplementary file 1 for "Spatially informed clustering, integration, and deconvolution of spatial transcriptomics with GraphST"

**Comparing the clustering performance of GraphST and existing methods on the mouse brain 10x Visium data.**

In this comparison, we tested GraphST and existing methods on a 10x Visium dataset of mouse brain anterior tissue, with 2,695 spots and 32,285 genes. This dataset has more complex tissue structures than the DLPFC dataset, posing a greater challenge. We manually annotated the spatial domains of this dataset using the Allen mouse brain reference atlas and set the number of clusters to 52 (Supplementary Figure S2). Similar to the DLPFC example, most of Seurat's clusters were fragmented, as were Giotto's and conST's. In the cortex region, all methods captured the layered structure with varying degrees of accuracy. Giotto, BayesSpace, SpaceFlow, and STAGATE produced more layers than expected based on the physiological annotation, while SpaGCN produced an abnormally thick cluster 1. In GraphST's output, the thickness of the different layers was closer to the annotation. The olfactory bulb also anatomically possessed a layered outer structure. Among the tested methods, SpaGCN was unable to capture this. In contrast, Giotto, BayesSpace, SpaceFlow, STAGATE, and GraphST captured more layers, with STAGATE and GraphST's results being the most similar. The computed clusters significantly overlapped across the methods, especially between BayesSpace, STAGATE, and GraphST. The caudoputamen (CPu) was another prominent structure that was divided into sub-regions. BayesSpace and STAGATE depicted it as two halves while GraphST captured a more layered structure. Overall, the results suggested that GraphST was more sensitive to the transcriptomic differences within the substructures.

**Comparing the deconvolution performance of GraphST and cell2location on the mouse brain 10x Visium data.**

We tested GraphST and cell2location for cell type deconvolution with the mouse brain 10x Visium data. A mouse scRNA-seq dataset encompassing the brain cortex and hippocampus was used as reference for deconvolution. The scRNA-seq data contains the single-cell transcriptomes of 1.1 million cells with 22,764 genes captured. The scRNA-seq data was annotated by the Allen Brain Map team to give 42 cell types. To reduce the computational burden, we downsampled the scRNA-seq data to 10%. We tested two approaches, random downsampling while retaining the same number of cell types, and proportional downsampling according to cell types, and found no performance difference. We first extracted the overlapping 1,099 genes in the scRNA-seq and ST data for training GraphST to map the single cell transcriptomes onto the spatial map. We then examined the mapped cell types found in the cerebral cortex by visualizing the scRNA-seq data cell types mapped onto specific spatial spots (Supplementary Figure S9). Both GraphST and cell2location correctly mapped the relevant cell types in the scRNA-seq data onto the ST positions of each cerebral cortex layer with GraphST's mapping showing sharper edges across all layers. In contrast, cell2location's mappings were more diffused, especially for the L4 RSP-ACA and L5 IT CTX cell types. GraphST and cell2location also mapped oligodendrocytes to the thalamus and mid brain with GraphST's mapping showing higher densities and sharper boundaries. The full results with all 42 cell types are given in Supplementary Figures S10 and S11 for GraphST and cell2location, respectively.

**GraphST's technical novelties and advantages over existing graph contrastive learning**

Existing methods conST and SpaceFlow also adopted graph contrastive learning similar to DGI for spatial transcriptomics analysis, but they were mainly developed for spatial clustering only. In addition to spatial clustering, GraphST can be also applied to two other important ST data analysis tasks, multi-sample integration and cell type deconvolution of ST. GraphST comprises three modules with different network architectures tailored for each of the three tasks respectively.

Even for the spatial clustering task, there are major technical differences and performance advantages when comparing GraphST to conST and SpaceFlow. Briefly, GraphST is different from DGI, conST, and SpaceFlow in three aspects: A) definition of positive/negative pairs, B) objective function and contrastive loss, and C) training procedure. These differences enable GraphST to outperform the other methods in the spatial clustering task. Furthermore, we have conducted several ablation studies to confirm that each of these differences improves effective integration of gene expression and spatial context to obtain informative and discriminative spot representations.

A) GraphST's contrastive learning is different from DGI, conST, and SpaceFlow in terms of their definition of positive/negative pairs. DGI, conST, and SpaceFlow construct positive/negative pairs by pairing each spot embedding $h_i/h_i'$ from the original/corrupted graph with a global summary vector $s_{global}$ (i.e., the mean of all spots' embeddings). Therefore, the spot embedding learned by DGI, conST, and SpaceFlow captures more of the global structure information but less spot-specific local neighbourhood information. Such contrastive learning may result in feature overfitting and reduced spot-to-spot variability. To deal with this issue, GraphST improves over DGI's contrastive learning by re-defining the positive/negative pairs. Specifically, motivated by the assumption that different spots in a tissue sample have different local spatial contexts, we define positive/negative pairs by pairing each spot embedding $h_i/h_i'$ with its local summary vector $s_{local}$ (i.e., the mean of its one-hop neighbouring spots' embeddings) instead of the global summary vector. With the local summary vector, the model can better preserve local context information and spot-to-spot variability. We demonstrate the effectiveness of local context with an ablation study describe below and the results are shown in Supplementary Figure S14A.

B) GraphST is also different from SpaceFlow, conST, and DGI in terms of the objective function and constructive loss formulations. The objective function of GraphST includes contrastive loss and reconstruction loss, while DGI's objective function includes only contrastive loss. The objective function of SpaceFlow includes contrastive loss and a spatial consistency penalty term. Addition of the penalty term helps SpaceFlow bring spatially adjacent spots closer in the latent embedding. However, the lack of reconstruction loss in DGI and SpaceFlow may lead to insufficient preservation of the original gene expression information. In contrast, GraphST adds reconstruction loss to its objection function to ensure that the latent embedding preserves the original gene expression information effectively. Furthermore, the contrastive loss functions are also different between GraphST, DGI, and SpaceFlow. GraphST adopts symmetric contrastive loss for model training while conST and SpaceFlow use single contrastive loss like DGI. Symmetric contrastive loss can help stabilize the model and learn a better representation as illustrated in Supplementary Figure S14B and S14E.

C) Lastly, although conST's objective function also contains contrastive loss and reconstruction loss like GraphST, GraphST's training procedure is different from that of conST. conST splits the training into pre-training and major training stages, where the model is trained with reconstruction loss in the pre-training stage and contrastive loss in the major training stage. This two-stage training procedure lacks mutual constraints on the contrastive and reconstructive loss, thus may fail to identify the optimal combination of the two loss functions. In contrast, GraphST trains the model in a single step by jointly optimizing the reconstruction and contrastive losses. During this training, GraphST can adaptively adjust the contributions of the different loss functions to achieve better representation learning.

**Ablation study**

For the spatial clustering module of GraphST, we define positive/negative pairs using local context instead of global context and employ symmetric contrastive loss instead of single contrastive loss. For spatial clustering, we use the reconstructed gene expression instead of the latent representation. Here we conducted ablation studies to evaluate their contributions to GraphST's performance.

Firstly, to evaluate the effectiveness of local context over global context, we conducted an ablation study by comparing GraphST with a variant that uses a global summary vector instead of local summary vectors. We ran GraphST and the variant on the 12 DLPFC slices and evaluated their performance using their median ARI scores. Supplementary Figure S14A shows that GraphST outperformed the variant with a significantly higher median ARI score of 0.60 than the variant (0.51). This study demonstrated that local context does help GraphST perform better than with the global context.

Secondly, to demonstrate the advantage of symmetric contrastive loss over single contrastive loss, we conducted an ablation study to compare GraphST with a variant that does not use contrastive corrupted loss. We tested GraphST and this variant on the 12 DLPFC samples and evaluated their performance with the ARI metric. Supplementary Figure S14B shows that GraphST achieved much better performance than the variant, illustrating that the contrastive corrupted loss contributes to better embedding learning. Furthermore, when employing contrastive learning with only a single contrastive loss, we found that the loss curve was unstable during model training (Supplementary Figure S14E). Motivated by the fact that the original and corrupted graphs are structurally identical, we added the symmetric (corrupted) contrastive loss function to make the model training more stable and robust. As we can see in Supplementary Figure S14E, the model-training curve was stabilized with the use of symmetric contrastive loss.

Thirdly, to demonstrate the contribution of self-supervised contrastive learning, we conducted an ablation study by comparing GraphST with a variant of GraphST without contrastive loss on the DLPFC dataset. Without contrastive loss, the performance of GraphST was significantly reduced by 15% (Supplementary Figure S14C), indicating that contrastive loss contributes to the performance improvement of our GraphST model.

Finally, we compared the clustering performance of using the latent representation and the reconstructed expression on the 12 DLPFC samples. The results (Supplementary Figure S14D) show that GraphST achieved a much higher median ARI score when using the reconstructed expression for clustering than the latent representation, suggesting that the former contains more useful information than the latter.

**Comparing Leiden, Louvain, and mclust clustering on GraphST outputs**

In GraphST, we chose mclust as the default clustering method because our assessment showed that mclust performs better than Leiden and Louvain in most cases. Here, we compared the clustering results on the 12 DLPFC samples using Leiden, Louvain, and mclust (Figure S15A). mclust consistently outperformed Leiden and Louvain on all 12 samples in terms of the ARI metric, with a much higher median ARI score (Figure S15B). Visually, the clusters identified by mclust are more continuous. Nevertheless, we also include Leiden and Louvain in GraphST as alternative clustering methods.

### Supplementary Figures

**Figure S1.** Manual annotations and comparison of spatial domains identified by Seurat, Giotto, SpaGCN, SpaceFlow, conST, BayesSpace, STAGATE, and GraphST on the 12 slices of DLPFC dataset.

**Figure S2.** Comparison of spatial domains identified by Seurat, Giotto, SpaGCN, SpaceFlow, conST, BayesSpace, STAGATE, and GraphST on the mouse brain anterior dataset. A. H&E image and manual annotation. B. Visualization of spatial domains identified by different methods.

**Figure S3.** Spatial clustering on the E9.5 mouse embryo Stereo-seq dataset. A. Tissue domain annotations obtained from the original Stereo-seq study. B. Spatial domains identified by GraphST. C. Visualization of spatial domains identified by GraphST. D. Visualization of marker genes supporting the identified domains.

**Figure S4.** Spatial clustering on the E14.5 mouse embryo Stereo-seq dataset. A. Tissue domain annotations obtained from the original Stereo-seq study. B. Spatial domains identified by GraphST when specified for 16 clusters. C. Visualization of spatial domains identified by the original Stereo-seq study and GraphST with 16 clusters. D. Spatial domains identified by GraphST when specified for 20 clusters. E. Gene set enrichment analysis on the cluster osteoblasts.

**Figure S5.** GraphST predicted spatial distributions of all 34 cell types in the human lymph node dataset.

**Figure S6.** cell2location predicted spatial distributions of all 34 cell types in the human lymph node dataset.

**Figure S7.** GraphST predicted spatial distributions of all 33 cell types in the 151673 slice of the DLPFC dataset.

**Figure S8.** cell2location predicted spatial distributions of all 33 cell types in the 151673 slice of the DLPFC dataset.

**Figure S9.** Comparison between cell2location and GraphST of spatial distributions of selected cell types in the mouse brain anterior dataset.

**Figure S10.** GraphST predicted spatial distributions of all 42 cell types in the mouse brain anterior dataset.

**Figure S11.** cell2location predicted spatial distributions of all 42 cell types in the mouse brain anterior dataset.

**Figure S12.** Comparison of spatial domains identified by Seurat, Giotto, SpaGCN, SpaceFlow, conST, BayesSpace, STAGATE, and GraphST on the human breast cancer data, and visualization of predicted spatial distribution of sample types. A. H&E image and manual annotation. B. Visualization of spatial domains identified by different methods. C. Violin plot of predicted probability of adjacent normal and solid tumor domains.

**Figure S13.** Spatial expression distribution of reported breast cancer markers.

**Figure S14.** Ablation study and comparison of training loss curves. A. Comparison analysis between GraphST and its variant using global context instead of local context. B. Comparison analysis between GraphST and its variant, i.e., GraphST without contrastive corrupted loss. C. Comparison analysis between GraphST and its variant, i.e., GraphST without contrastive loss. D. Comparison analysis between GraphST and its variant using latent representation for clustering instead of reconstructed expression. E. Training loss curves with and without contrastive corrupted loss on four DLPFC samples.

**Figure S15.** Comparison analysis between Leiden, Louvain, and mclust with the output of GraphST as input with the DLPFC dataset. A. Visualization of clustering results from Leiden, Louvain, and mclust on 12 DLPFC slices. B. Boxplots of clustering accuracy of Leiden, Louvain, and mclust on 12 DLPFC slices.

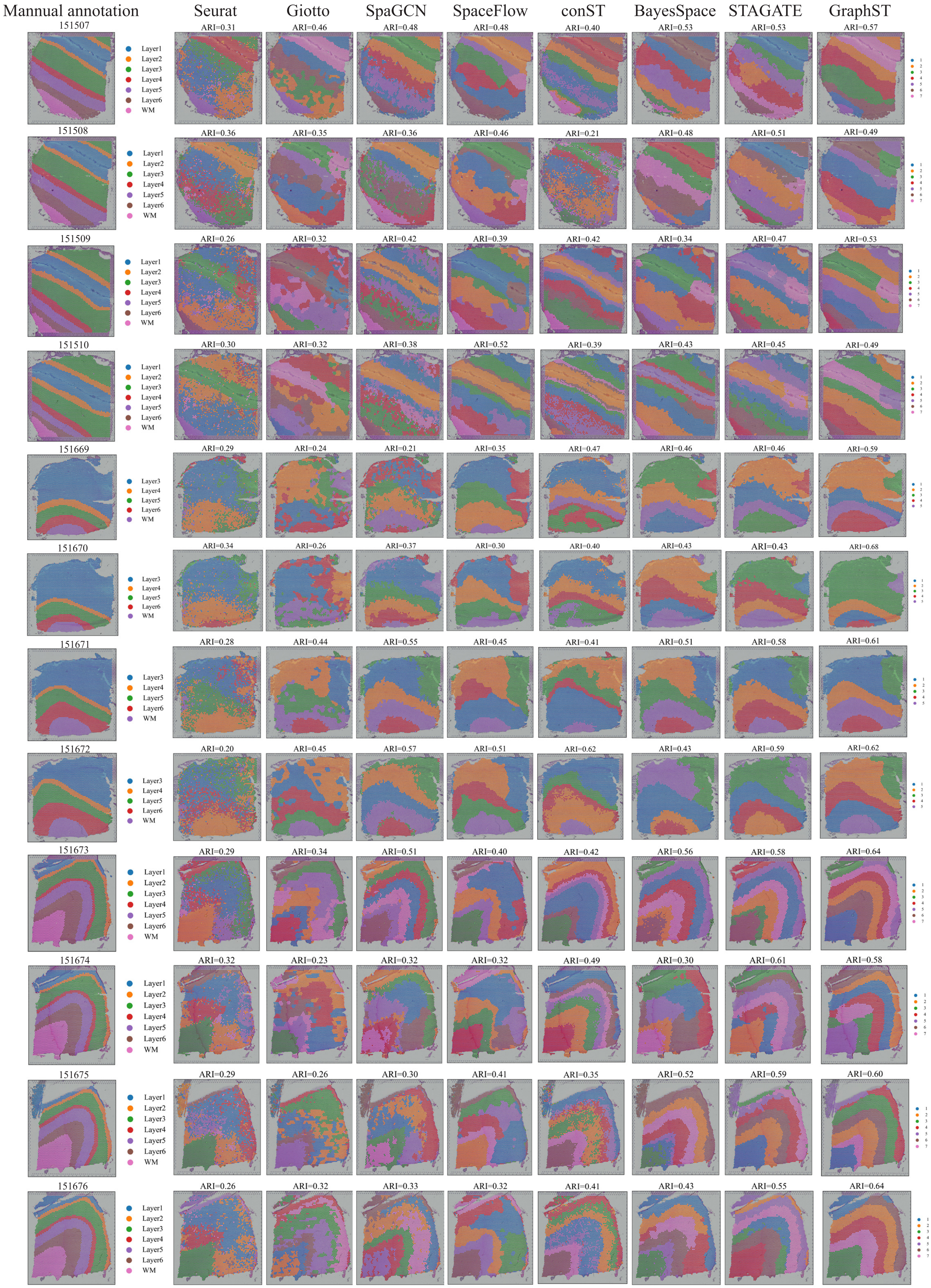


Figure S1. Manual annotations and comparison of spatial domains identified by Seurat, Giotto, SpaGCN, SpaceFlow, conST, BayesSpace, STAGATE, and GraphST on the 12 slices of DLPFC dataset.

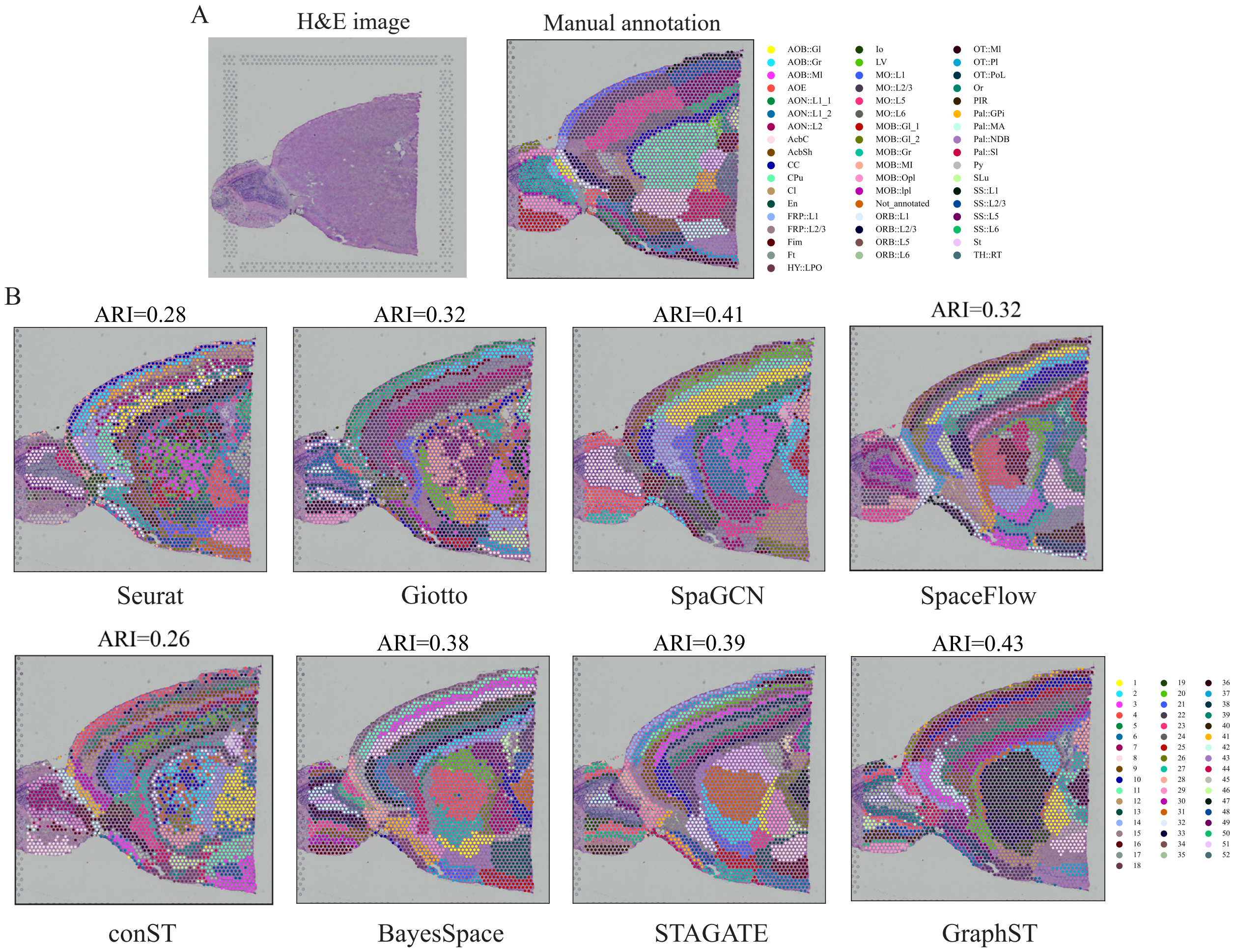


**Figure S2. Comparison of spatial domains identified by Seurat, Giotto, SpaGCN, SpaceFlow, conST, BayesSpace, STAGATE, and GraphST on the mouse brain anterior dataset.** A. H&E image and manual annotation. B. Visualization of spatial domains identified by different methods.

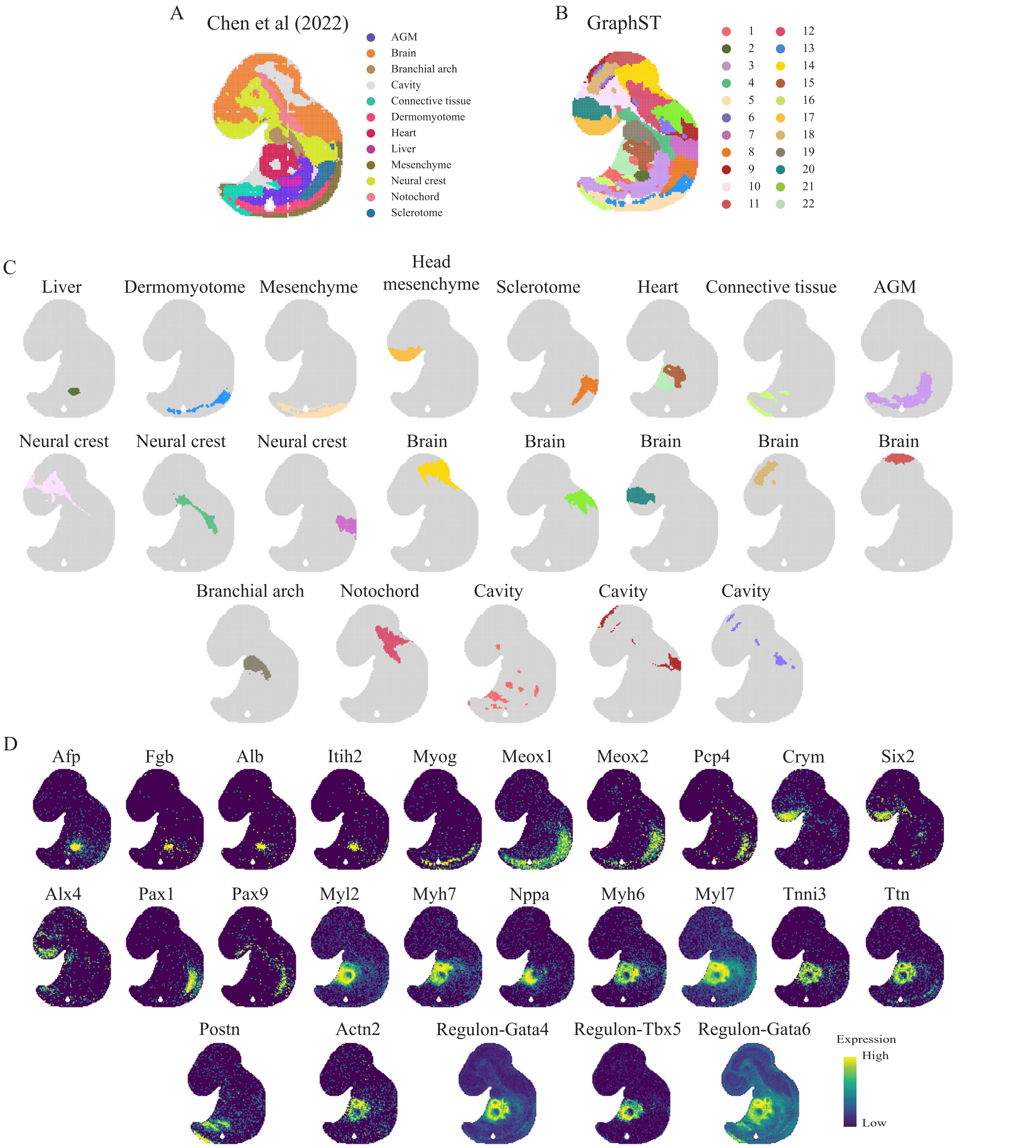


**Figure S3. Spatial clustering on the E9.5 mouse embryo Stereo-seq dataset. A.** Tissue domain annotations obtained from the original Stereo-seq study. **B.** Spatial domains identified by GraphST. **C.** Visualization of spatial domains identified by GraphST. **D.** Visualization of marker genes supporting the identified domains.

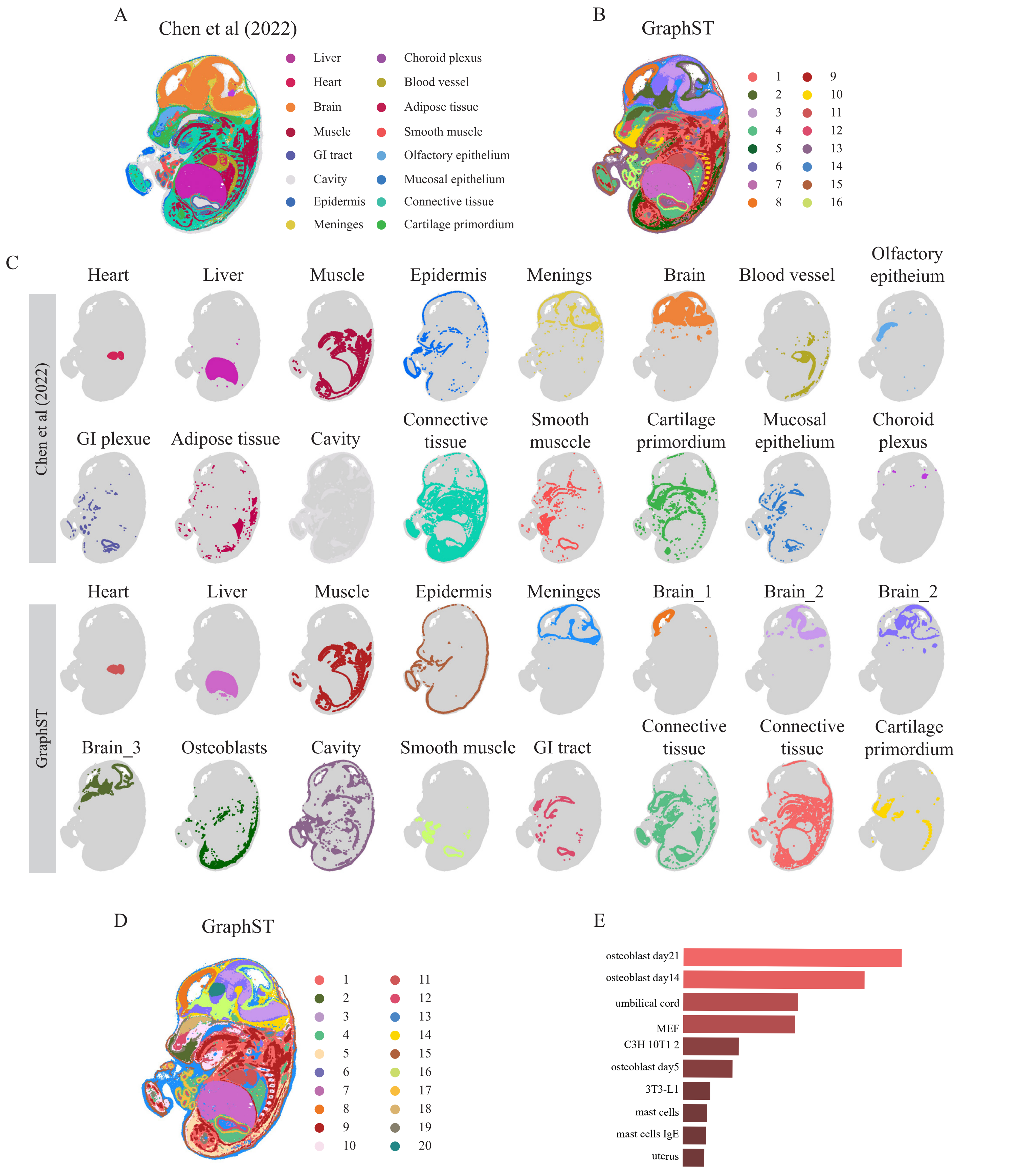


**Figure S4. Spatial clustering on the E14.5 mouse embryo Stereo-seq dataset. A.** Tissue domain annotations obtained from the original Stereo-seq study. **B.** Spatial domains identified by GraphST when specified for 16 clusters. **C.** Visualization of spatial domains identified by the original Stereo-seq study and GraphST with 16 clusters. **D.** Spatial domains identified by GraphST when specified for 20 clusters. **E.** Gene set enrichment analysis on the cluster osteoblasts.

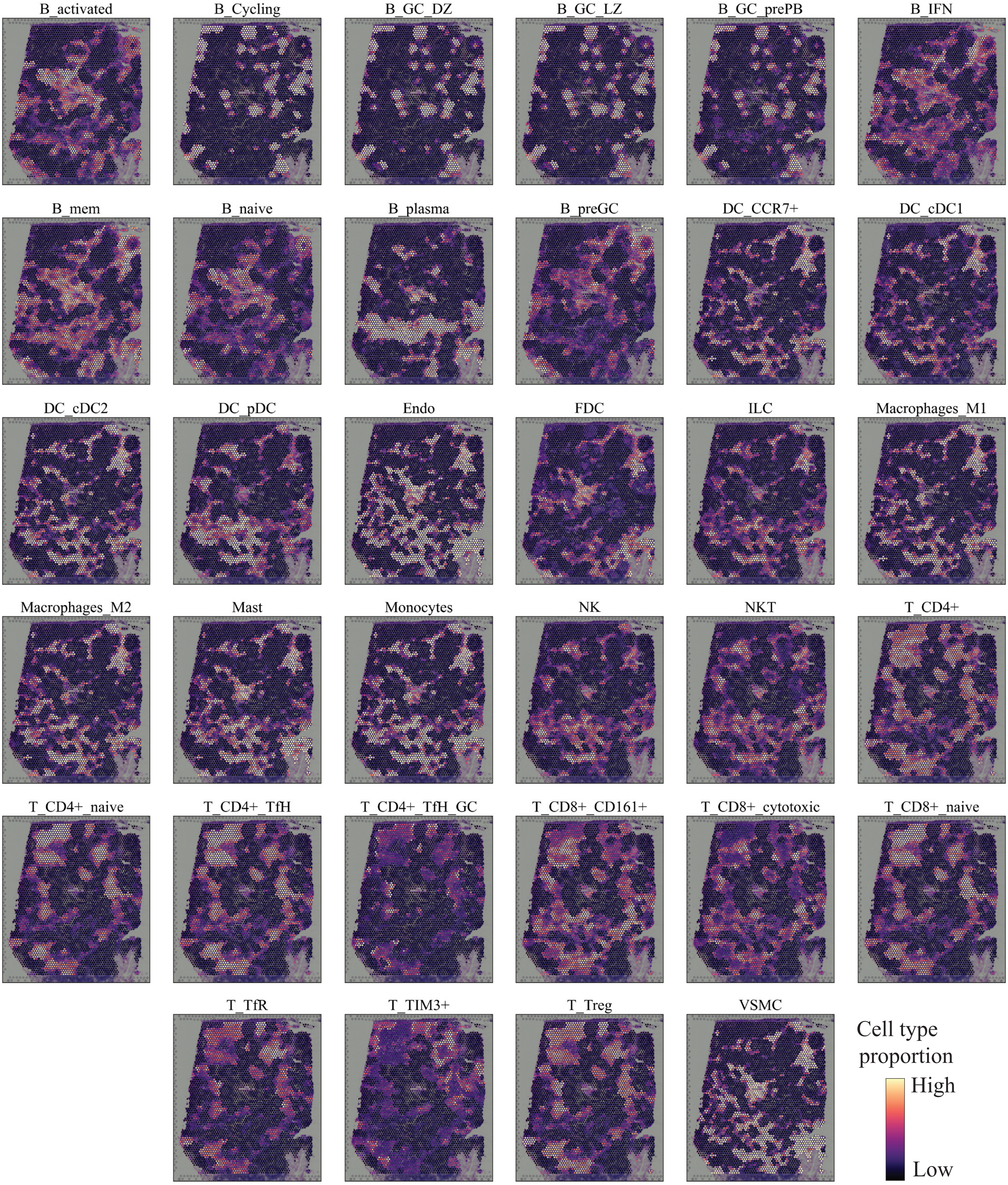


Figure S5. GraphST predicted spatial distributions of all 34 cell types in the human lymph node dataset.

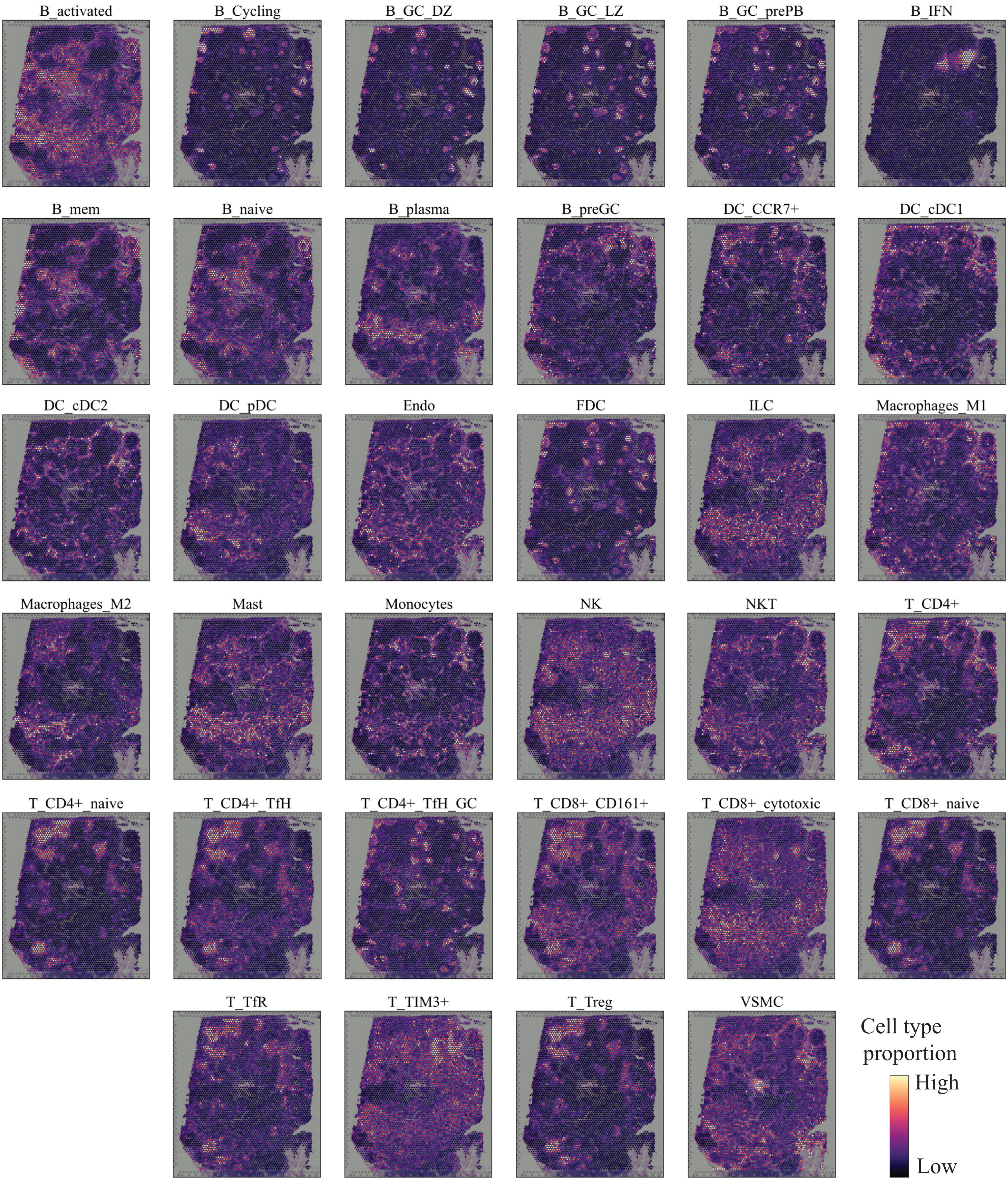


Figure S6. cell2location predicted spatial distributions of all 34 cell types in the human lymph node dataset.

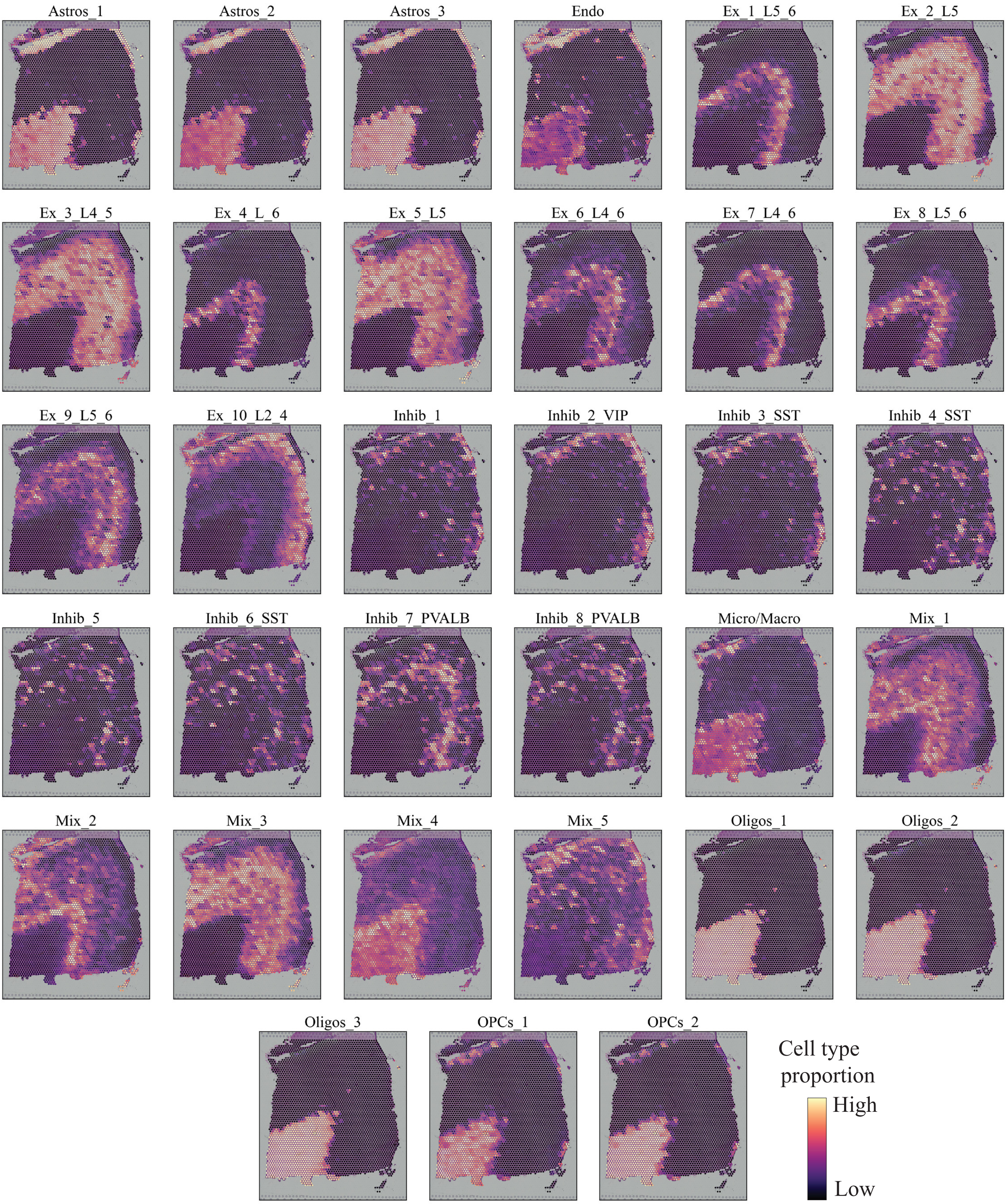


Figure S7. GraphST predicted spatial distributions of all 33 cell types in the 151673 slice of the DLPFC dataset.

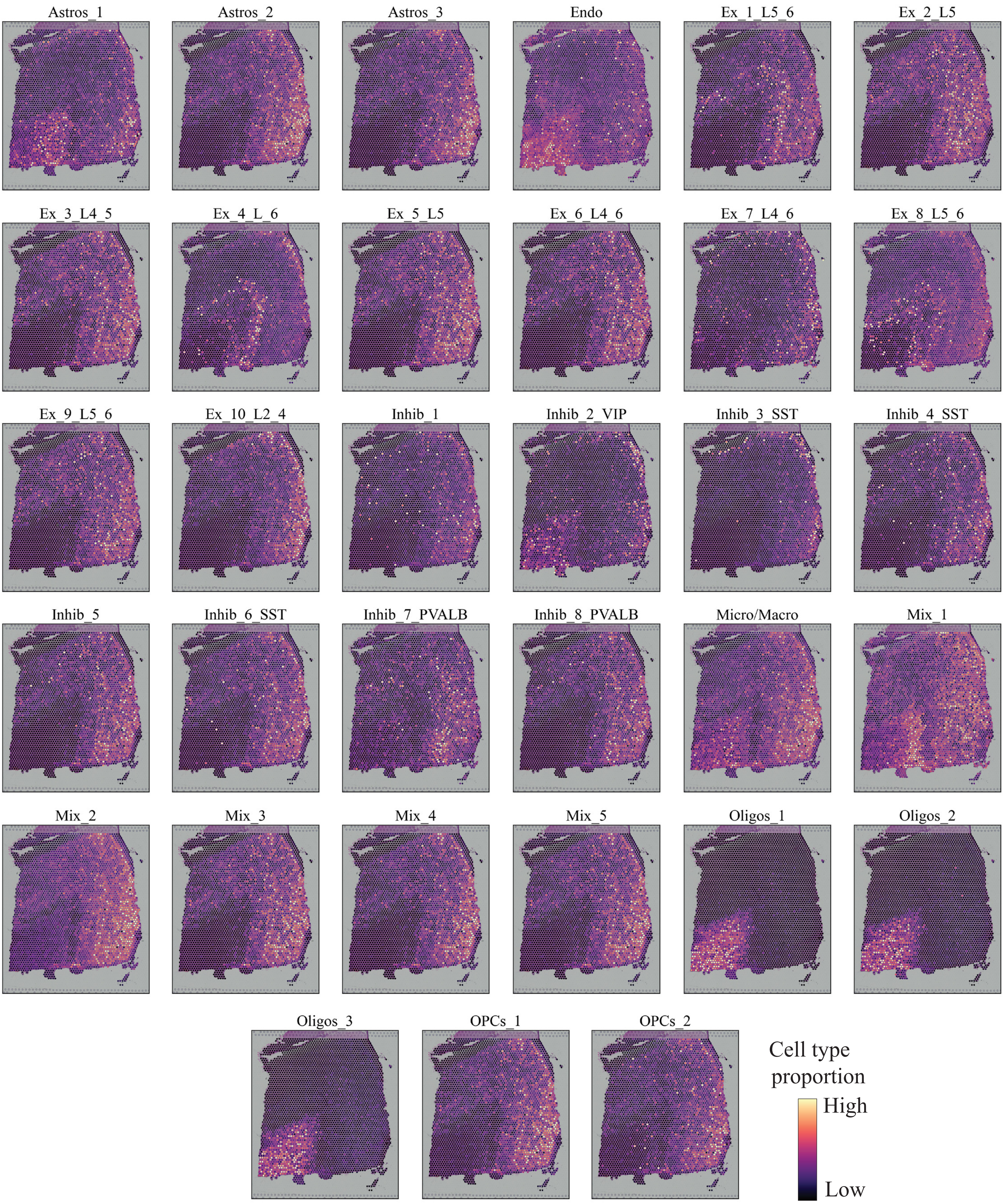


Figure S8. cell2location predicted spatial distributions of all 33 cell types in the 151673 slice of the DLPFC dataset.

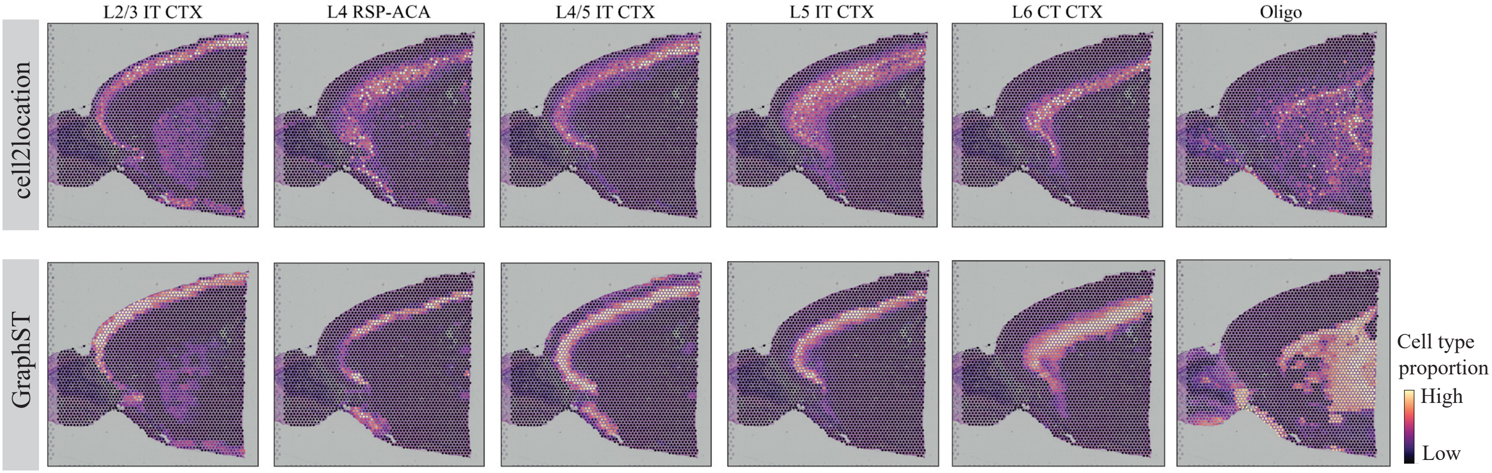


Figure S9. Comparison between cell2location and GraphST of spatial distributions of selected cell types in the mouse brain anterior dataset.

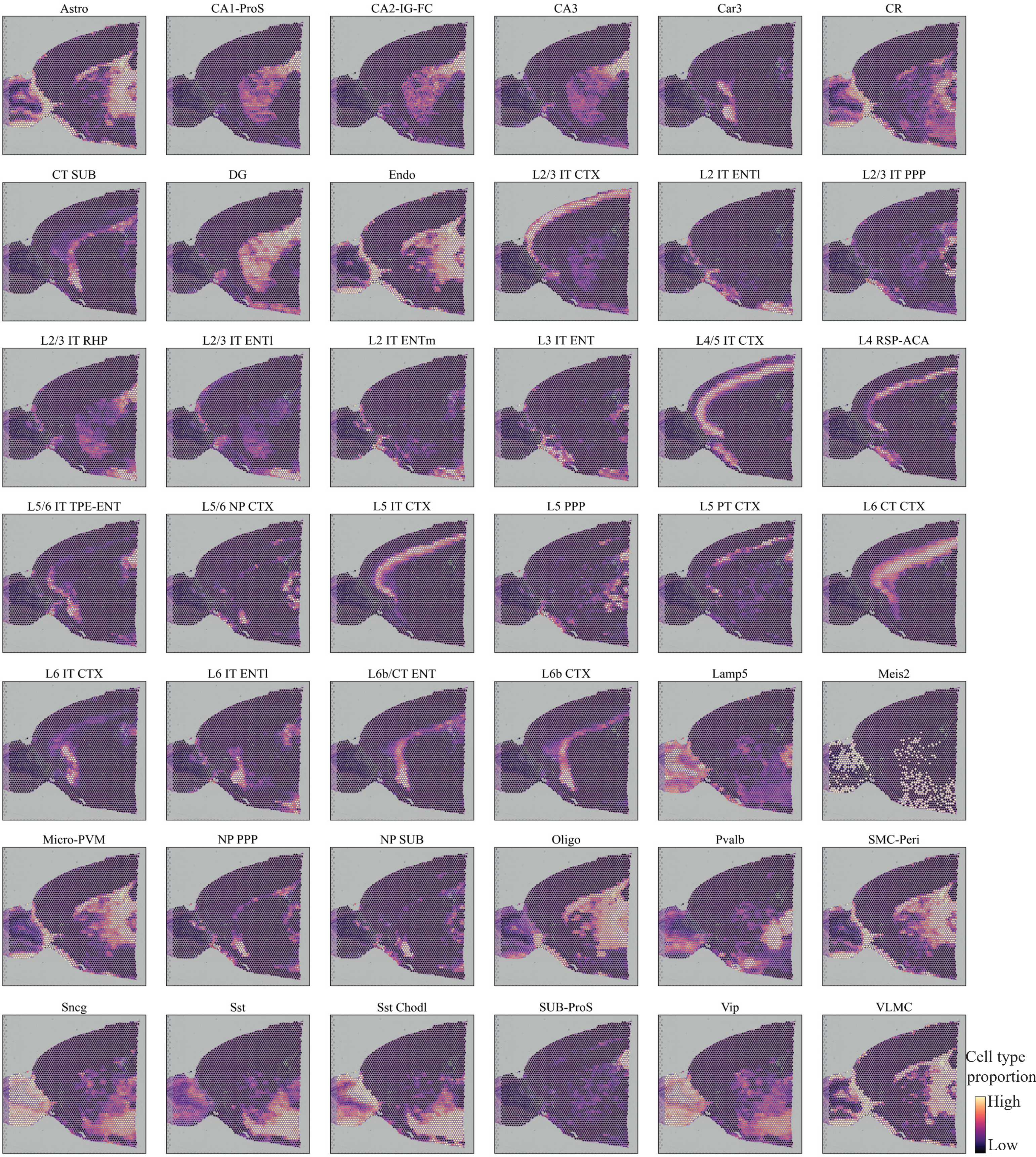


Figure S10. GraphST predicted spatial distributions of all 42 cell types in the mouse brain anterior dataset.

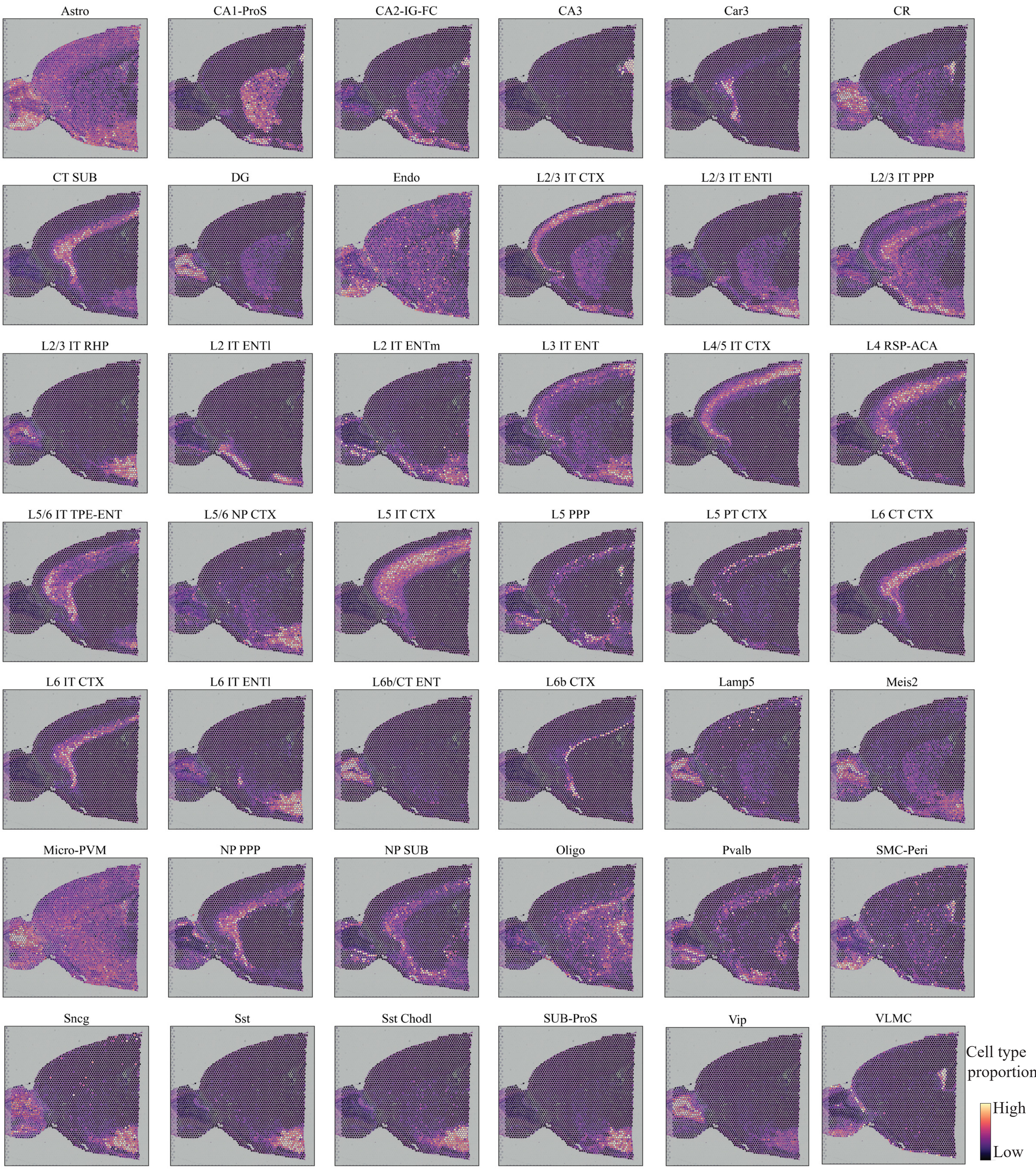


Figure S11. cell2location predicted spatial distributions of all 42 cell types in the mouse brain anterior dataset.

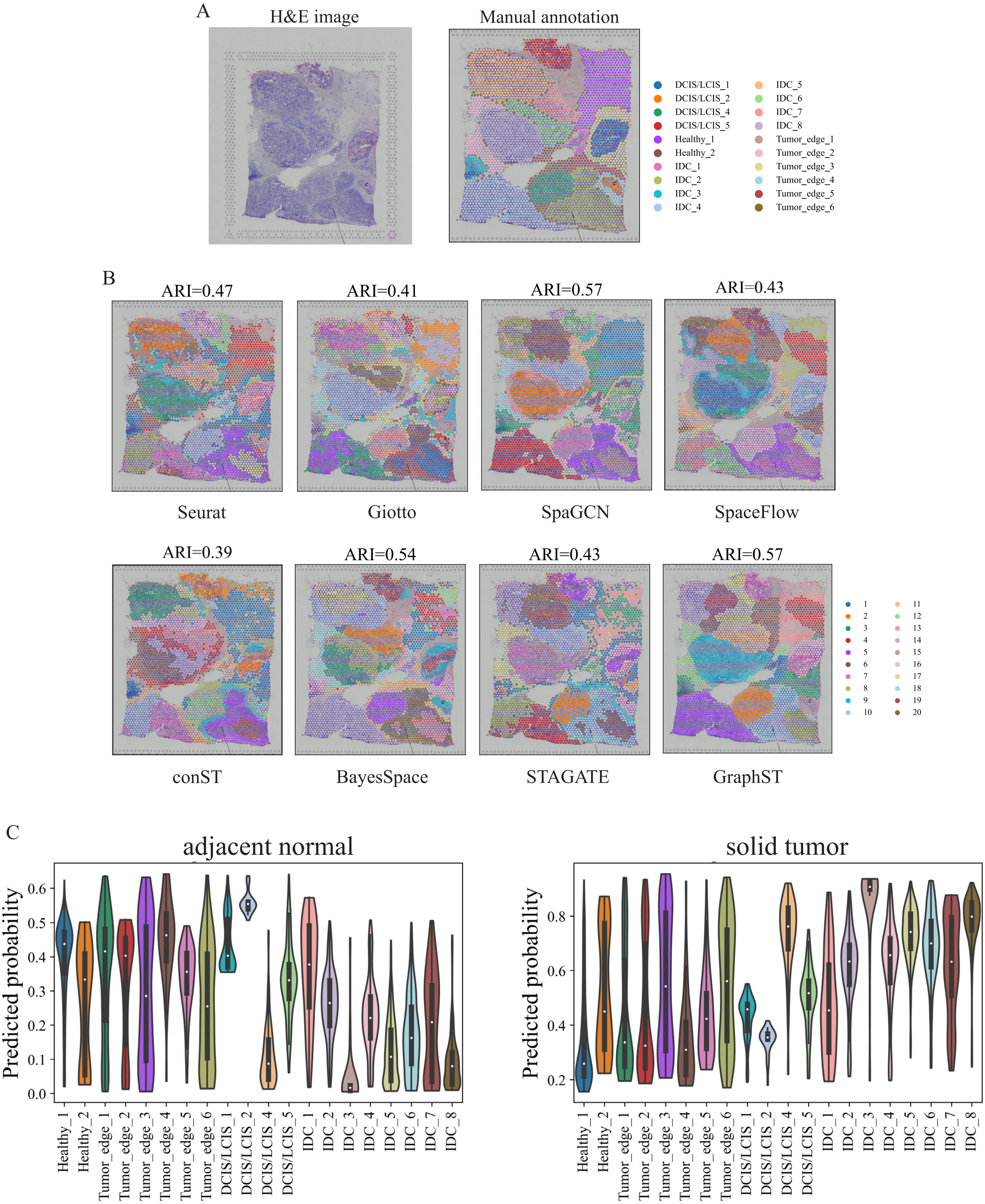


**Figure S12. Comparison of spatial domains identified by Seurat, Giotto, SpaGCN, SpaceFlow, conST, BayesSpace, STAGATE, and GraphST on the human breast cancer data, and visualization of predicted spatial distribution of sample types. A.** H&E image and manual annotation. **B.** Visualization of spatial domains identified by different methods. **C.** Violin plot of predicted probability of adjacent normal and solid tumor domains.

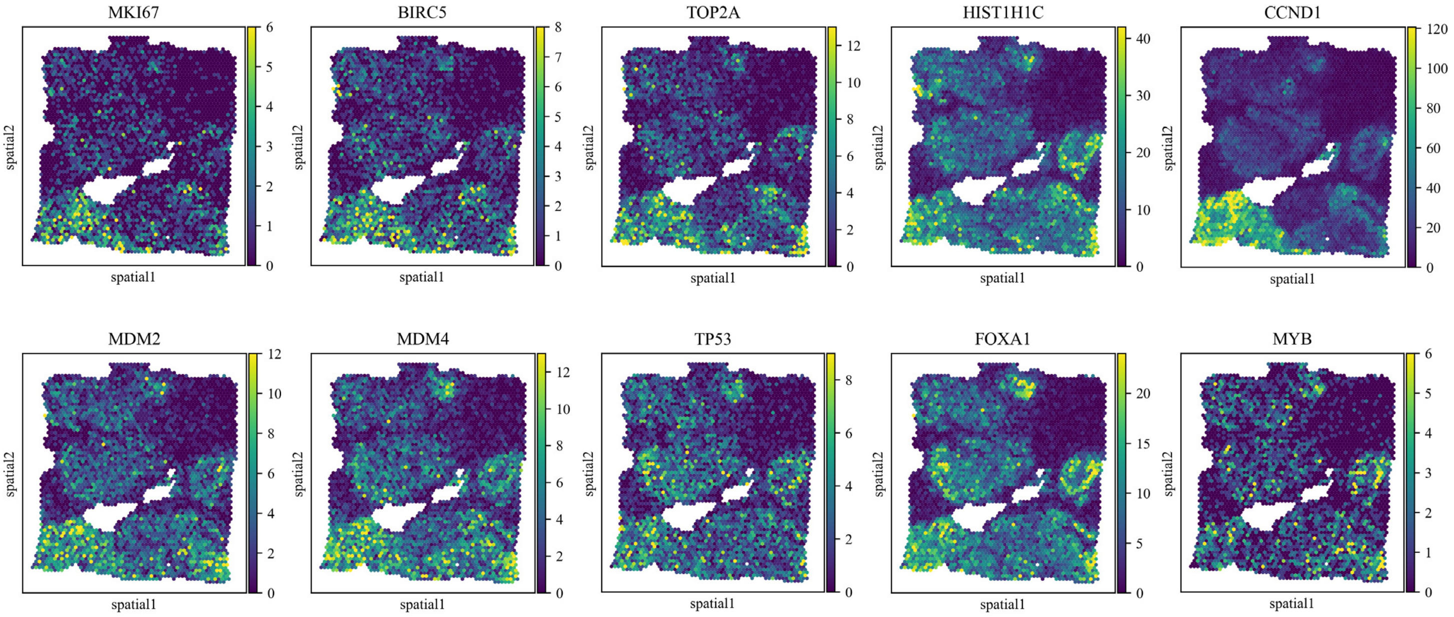


Figure S13. Spatial expression distribution of reported breast cancer markers.

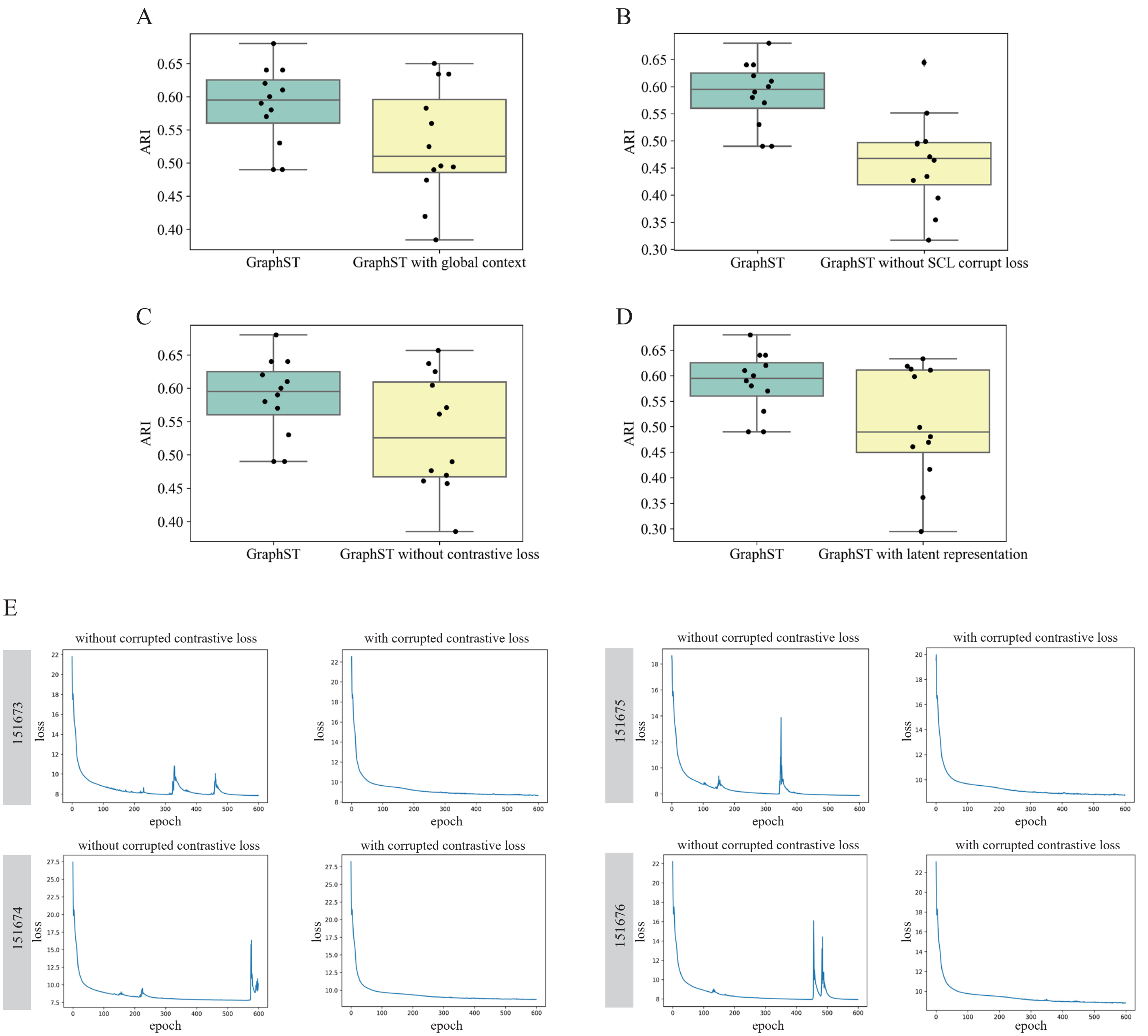


**Figure S14. Ablation study and comparison of training loss curves. A.** Comparison analysis between GraphST and its variant using global context instead of local context. **B.** Comparison analysis between GraphST and its variant, i.e., GraphST without contrastive corrupted loss. **C**. Comparison analysis between GraphST and its variant, i.e., GraphST without contrastive loss. **D**. Comparison analysis between GraphST and its variant using latent representation for clustering instead of reconstructed expression. **E**. Training loss curves with and without contrastive corrupted loss on four DLPFC samples.

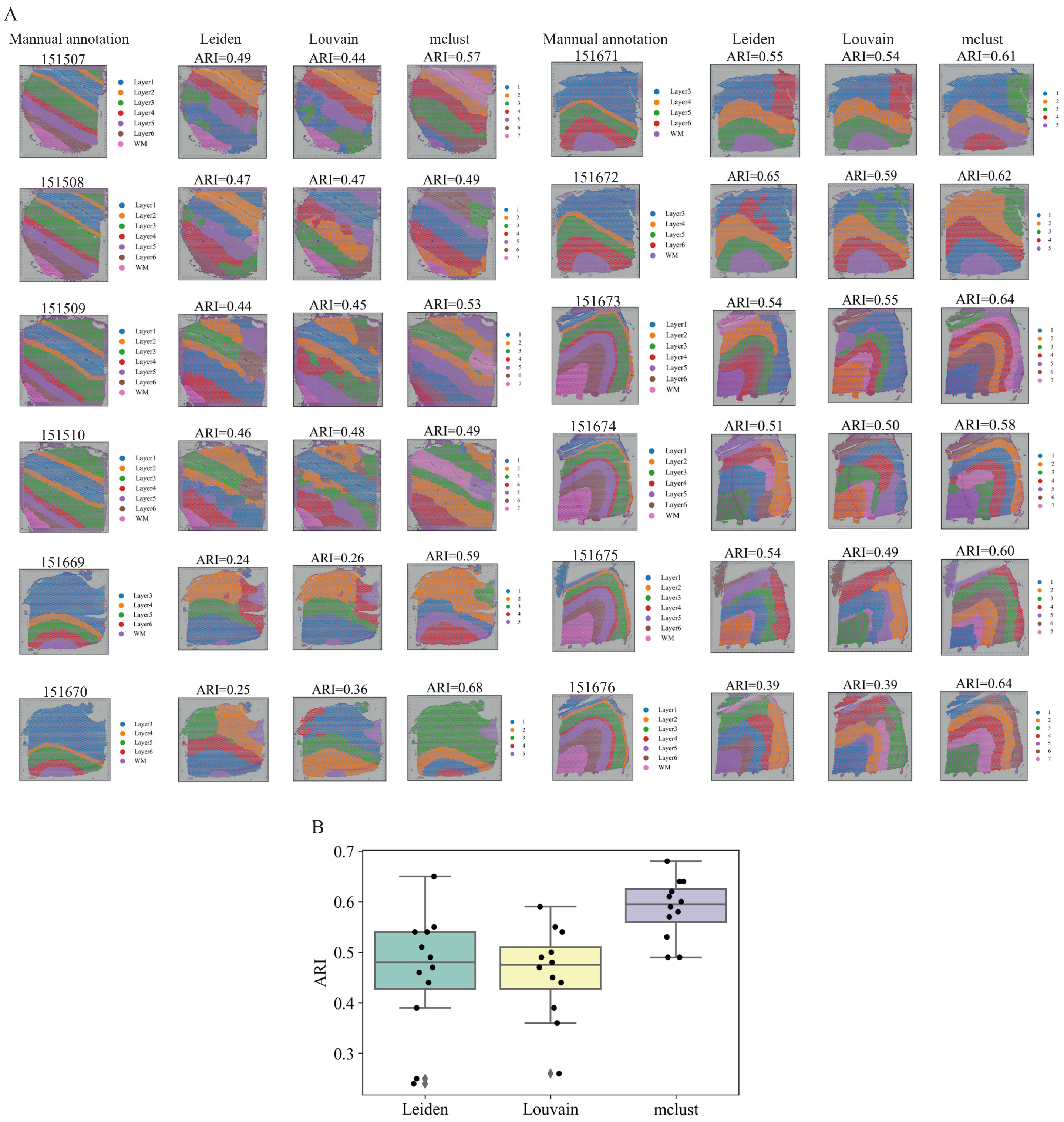


**Figure S15. Comparison analysis between Leiden, Louvain, and mclust with the output of GraphST as input with the DLPFC dataset. A.** Visualization of clustering results from Leiden, Louvain, and mclust on 12 DLPFC slices. **B.** Boxplots of clustering accuracy of Leiden, Louvain, and mclust on 12 DLPFC slices.
